# Escape from X-inactivation in twins exhibits intra- and inter-individual variability across tissues and is heritable

**DOI:** 10.1101/2021.10.15.463586

**Authors:** A. Zito, A.L. Roberts, A. Visconti, N. Rossi, R. Andres-Ejarque, S. Nardone, J.E.S. Moustafa, M. Falchi, K.S. Small

## Abstract

X-chromosome inactivation (XCI) silences one X-chromosome in female cells to balance sex-differences in X-dosage. A subset of X-linked genes escape XCI, but the extent to which this phenomenon occurs and how it varies across tissues and in a population is as yet unclear. In order to characterize the incidence and variability of escape across individuals and tissues, we conducted a large scale transcriptomic study of XCI escape in adipose, skin, lymphoblastoid cell lines (LCLs) and immune cells in 248 twins drawn from a healthy population cohort. We identify 159 X-linked genes with detectable escape, of which 54 genes, including 19 lncRNAs, were not previously known to escape XCI. Across tissues we find a range of tissue-specificity, with 11% of genes escaping XCI constitutively across tissues and 24% demonstrating tissue-restricted escape, including genes with cell-type specific escape between immune cell types (B, T-CD4^+^, T-CD8^+^ and NK cells) of the same individual. Escape genes interact with autosomal-encoded proteins and are involved in varied biological processes such as gene regulation. We find substantial variability in escape between individuals. 49% of genes show inter-individual variability in escape, indicating escape from XCI is an under-appreciated source of gene expression differences. We utilized twin models to investigate the role of genetics in variable escape. Overall, monozygotic (MZ) twin pairs share more similar escape than dizygotic twin pairs, indicating that genetic factors underlie differences in escape across individuals. However, we also identify instances of discordant XCI within MZ co-twin pairs, suggesting that environmental factors also influence escape. Thus, XCI escape may be shaped by an interplay of genetic factors with tissue- and cell type-specificity, and environment. These results illuminate an intricate phenotype whose characterization aids understanding the basis of variable trait expressivity in females.

## Introduction

The X chromosome is a paradigmatic genetic model^1^. It carries >1000 genes, representing >5% of the haploid human genome. It is differentially inherited between the sexes. The unequal X-linked transcriptional dosage between the sexes is partially compensated by random silencing of one X in each female somatic cell^2^. This process, known as X-chromosome inactivation (XCI), involves synergistic DNA-RNA-protein interactions that mediate heterochromatinization of the X designated for inactivation^1,3,4^ (known as “Barr Body”^5^). Non-coding RNAs play key roles in XCI. The master long non-coding (lnc) RNA *XIST* spreads in *cis* along the inactive X chromosome (Xi) and promotes a progressive epigenetic silencing^4,6,7^. However, XCI is incomplete, with over 15% of X-genes reported to escape silencing and exhibit expression from both parental alleles within a diploid cell^8,9^. Mary Lyon predicted that genes with Y-homologues (e.g. pseudo autosomal regions (PARs)), are naturally dosage compensated and thus expected to escape^10^. Today, most known escapees lack functional Y-homologues, thus being a potential source of sexual dimorphism^9,11^. Chromosome X is enriched for genes with immune- and neuro-modulatory functions^12,13^; changes in escape can thus underpin not only sexual dimorphism, but also phenotypic and disease risk variability across females^12,14,15^. Despite its biomedical relevance, the inter-individual variability of escape at population level, and across cells and tissues within an individual, has not been systematically examined. Furthermore, the extent to which genetics and environment influence the escape remains largely undefined.

Our current knowledge of XCI escape in humans largely rely on conventional studies of male/female expression ratio, human/mouse hybrid cells, and epigenetic marks^8,11,16,17^. In most females, the random nature of XCI results in expression of both X-linked alleles at a tissue level. This limits the ability to distinguish mono-from biallelic expression and thus identification of escape^9^. To circumvent the problem, strategies like single-cell analyses (e.g. scRNAseq) or sex comparison have been used to infer escape^9,11,18,19^. However, scRNAseq is infeasible for large cohorts and is limited to highly and consistently expressed genes due to allelic drop-out and transcriptional burst, which both can inflate monoallelic expression ratio. On the other hand, sex differences may not directly reflect the allelic expression ratio of X-genes in a tissue. These limitations can be circumvented by using tissue samples exhibiting skewed XCI patterns – a common event in the female population^20^ – which, as opposed to random XCI, enable detection and measurement of escape directly in skewed females (Note S1). This strategy has been employed, but either sample sizes were limited (e.g. single GTEx donor), or the study relied on arrays and other artificial biological models^8,11,21^.

Here, we characterize XCI escape using paired bulk RNAseq and high-coverage DNAseq data in a multi-tissue dataset sampled from 248 skewed female twins of the TwinsUK bioresource^22^. We investigate escape prevalence and variability across adipose and skin tissue, lymphoblastoid cell lines and purified immune cells (monocytes, B-cells, T-CD4^+^, T-CD8^+^ and NK cells), and individuals. We identify novel genes exhibiting tissue- and immune cell type-specific escape, and genes escaping XCI with high variability across tissues and individuals. We observe that escape varies across tissues and immune cells within an individual and across individuals, a phenomenon with high biomedical relevance. Using twins, we demonstrate that regulation of XCI escape has both heritable and environmental components, implying a complex interplay between genetic and non-genetic factors.

## RESULTS

### Escape from XCI is more prevalent in solid tissues and wider than currently known

We quantified escape in multiple tissues concurrently sampled from female twins of the TwinsUK cohort^22^. We determined XCI patterns using the gene-level *XIST* allele-specific expression (*XIST*_ASE_) from paired RNAseq and DNAseq data^7,23,24^. From over 2200 tissue samples interrogated, we obtained *XIST*_ASE_ calls for 522 LCLs, 101 whole-blood, 421 adipose, and 373 skin samples. In samples exhibiting skewed XCI (0.2≥*XIST*_ASE_≥0.8, Methods) including 166 LCLs (32%), 26 whole-blood (26%), 57 adipose (14%), and 64 skin (17%) samples, the levels of escape of each X-linked gene were measured using a metric - herein referred to as Escape Score or ‘EscScore’ - derived from the gene’s allelic fold-change adjusted for the sample’s XCI skew (Methods). EscScore values range from 0 (no escape, monoallelic expression) to 1 (full escape, equal expression from inactive (Xi) and active X (Xa)). We interrogated a total of 551 genes, of which 85% are protein-coding and 15% non-coding RNA genes. Based on a publicly available catalogue of XCI statuses^25^ (‘Balaton’s list’), our interrogated genes were categorized as XCI-silenced (n=326), fully or mostly escaping XCI (n=52), or variable escapees (n=23). Variable escape refers to genes whose escape is variable across cells, tissues, or individuals^8,11,25^. We also included a subset of 41 genes whose XCI status was reported as discordant across studies or undefined^25^. The summary statistics indicated that EscScore differs between different categories (Table S1). In all tissues, the EscScore of genes annotated as fully or mostly escaping XCI significantly differed from genes annotated as either silenced or variable escapees (Fig.1; Table S2), supporting the reliability of our escape metric to discriminate different XCI statuses.

**FIG.1.**
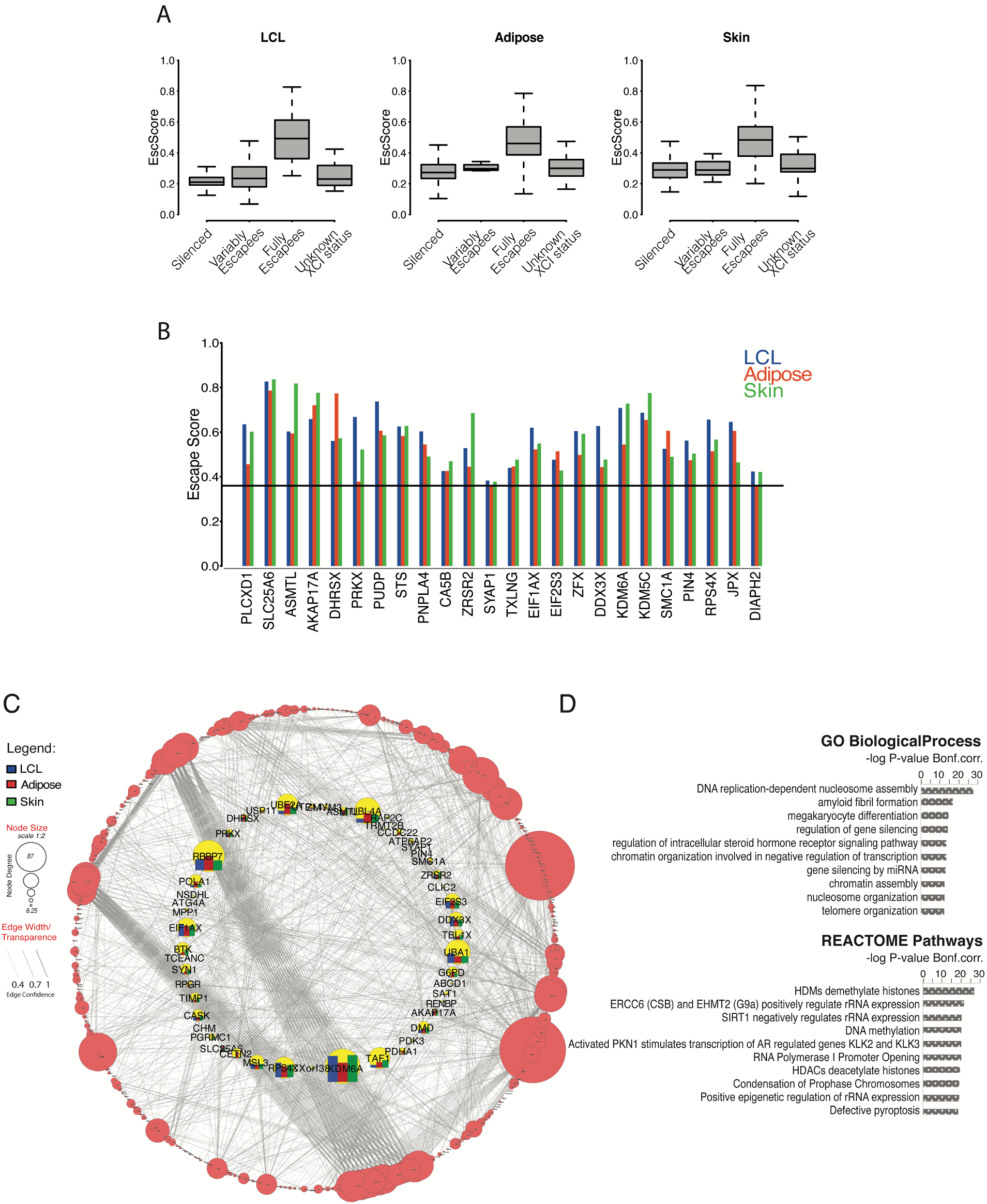
(A) EscScore distinguishes between silenced and escape genes. Boxplots show the distribution of median EscScore values (across skewed samples) of genes with different previously annotated XCI status^25^. (B) Genes exhibiting constitutive XCI escape in all three tissues. Barplots show the tissue-specific gene’s EscScore. Horizontal line denotes the EscScore cutoff to call XCI escape. (C-D) PPI, Gene Ontology (GO) and Pathway analysis. (C) PPI network between protein-coding X-genes escaping XCI in at least one tissue and a reference map of human proteome interactome^28^. The network inner circle shows protein-coding escape genes characterized within the proteome interactome. The network outer circle shows proteins from the reference proteome^28^ that have (i) at least 1 interaction with our escapee genes and (ii) edge confidence score ≥0.4. The size of each node reflects the number of interactions (edges). (D) GO and REACTOME Pathways analysis of PPI network. The top 10 Biological Processes and REACTOME terms are shown.

By defining the allele with the lower RNAseq count as the Xi allele while the other as Xa allele, we detected an average Xi/Xa expression ratio of 0.32, 0.36, and 0.37 across LCLs, adipose and skin samples, respectively, with median value of 0.36. This is consistent with the GTEx survey of XCI, which reported that the Xi expression is on average 33% of Xa expression^11^. Based on these observations, we classified genes with a median (across ≥3 skewed tissue samples) EscScore≥0.36 as escapees in that tissue, while genes with EscScore<0.36 as silenced. In line with current knowledge, most X-genes are subject to XCI in all tissues (84% in LCLs, 74% in adipose, 71% in skin). We observed a higher incidence of escape in solid tissues than LCLs, with 16%, 26% and 29% of genes escaping XCI in LCLs, adipose and skin tissues, respectively. Altogether, 159 X-genes exhibited escape in at least one tissue in our dataset (Table S3). Expectedly, PAR-linked genes escaped XCI. We used a hypergeometric test to assess overlap between the Balaton’s list^25^ and our list of escapees, and found significant overlap (N=50; *P*≤0.05). Among our escape calls, about 60 genes retain a Y-pseudogene or Y-homologue, and 13 are PAR-linked, supporting their escape. For a more comprehensive comparison, we merged the Balaton’s list of escapees with additional external lists of escapees from other studies including (i) the GTEx XCI survey^11^; (ii) Zhang *et al*.^26^; (iii) Katsir *et al*.^19^; (iv) Garieri et al.^18^. We found that 66% (N=105) of our escapees overlapped with this unified list of escapees (*P* ≤0.05), while the remaining (N=54) represent novel calls. We confirmed the escape status of *CLIC2* in skin, as identified in GTEx^11^, and also found it escapes in adipose in our data. Notably, 35% of our novel escape calls are annotated as lncRNA, while the remaining are protein-coding. We establish the first evidence that the lncRNA *AL683807*.*1* escapes XCI. *AL683807*.*1* is PAR1-linked, explaining its escape ability. We examined the chromosomal distribution of escape genes and confirmed a higher escape incidence on the short arm^27^ (Note S2; Fig.S1). Altogether, these data further support the suitability of our study design. We show that in our data, escape is more prevalent in adipose and skin than LCLs, and is wider than currently known.

### Escape from X-inactivation exhibits both constitutive and tissue-specific patterns

Presently, the extent to which tissue-specific escape occurs in humans is unclear. To study this, we calculated the gene’s median EscScore (across ≥3 tissue samples) as a measure for tissue-specific levels of escape. We found significant differences across tissues (Kruskal-Wallis (‘KW’) *P-value<*10^−10^; Fig.S2), suggesting tissue-specific components of escape. Using a subset of 215 genes with EscScore available in all tissues (Table S4), we identified 24 genes exhibiting escape in both LCLs and solid tissues (Fig.1B), suggesting constitutive escape. We observed that tissue-specific EscScore remained below 80% in most cases, in line with data showing that Xi/Xa expression ratio would not exceed 80%^11^. Notably, *PLCXD1, ASMTL, DHRSX, SLC25A6* and *AKAP17A* are PAR-linked. We show that *PUDP* and *PIN4*, whose escape status have remained so far unclear, show constitutive escape in all tissues. Among the constitutive escapees, there are the highly biomedically-relevant genes *DDX3X, KDM5C* and *KDM6A*, whose escape may contribute to lower cancer incidence in females than males^14^. We also observed that *PRKX, PUPD, DDX3X* and *JPX* each had significantly different escape between tissues (KW *P*≤0.05). Tissue-specificity of escape was further supported by identification of 51 genes exhibiting escape restricted to a single tissue. Notably, 18 of these, of which 3 non-coding RNAs and 15 protein-coding genes, are novel escape calls (Table S5).

We investigated whether the escapees may interact with other factors and be involved in biological processes. To address this, we selected genes exhibiting escape in at least one tissue and conducted protein-protein interaction network analysis using a recent human protein interactome as reference^28^. We found that protein-coding escape genes interact with other factors on a genome-wide scale (Fig.1C). Gene ontology analysis revealed that members of this proteome network are involved in distinct biological processes such as epigenetic regulation by chromatin assembly and nucleosome organization, and regulation of steroid hormone signaling. These data were supported by REACTOME pathway analysis which revealed multiple pathways for epigenetic control of genes, such as histone methylation and acetylation, and DNA methylation (Fig.1D, Table S6). Altogether, these data suggest that escape is shaped by an interplay of tissue-shared and tissue-specific factors, and participates in genome-wide interactions involved in varied biological processes.

### Escape from X-inactivation exhibits intra- and inter-individual variability

The extent to which escape varies across tissues within an individual is unclear. We investigated this phenomenon in a subset of 6 donors exhibiting skewed XCI in LCLs, adipose and skin. This strategy was employed in the GTEx survey using a single skewed female donor^11^. Within each donor, we examined all genes with available EscScore. We identified genes exhibiting escape (EscScore≥0.36 in a tissue within a donor) in 1, 2 or 3 tissues and found that their prevalence varied between donors (Fig.2A; Table S7). Occurrence of genes escaping XCI in all tissues in multiple donors suggests shared regulatory mechanisms across tissues and individuals. Genes exhibiting such a behaviour included the zinc-finger protein *ZFX* and the histone demethylase *KDM5C* which is linked to intellectual disability and autism^29,30^. We also observed genes escaping XCI in all tissues but in only 1 of the 6 donors. Examples are the leukaemia-protecting histone demethylase *KDM6A*^14,31^, *FMR1*, a gene linked to Fragile-X and learning disability^32^, and the Duchenne muscular dystrophy gene *DMD*^*33*^. We observed that *ASMTL* escaped XCI in all tissues in a donor, and the degree to which it escaped varied across tissues ranging from 0.6 to over 0.8. *ASMTL*’s behavior was also highlighted in GTEx^11^. Aside from these cases, most of the interrogated genes exhibited tissue-restricted escape, supporting the occurrence of tissue-specific factors exerting dominant effects.

**FIG.2.**
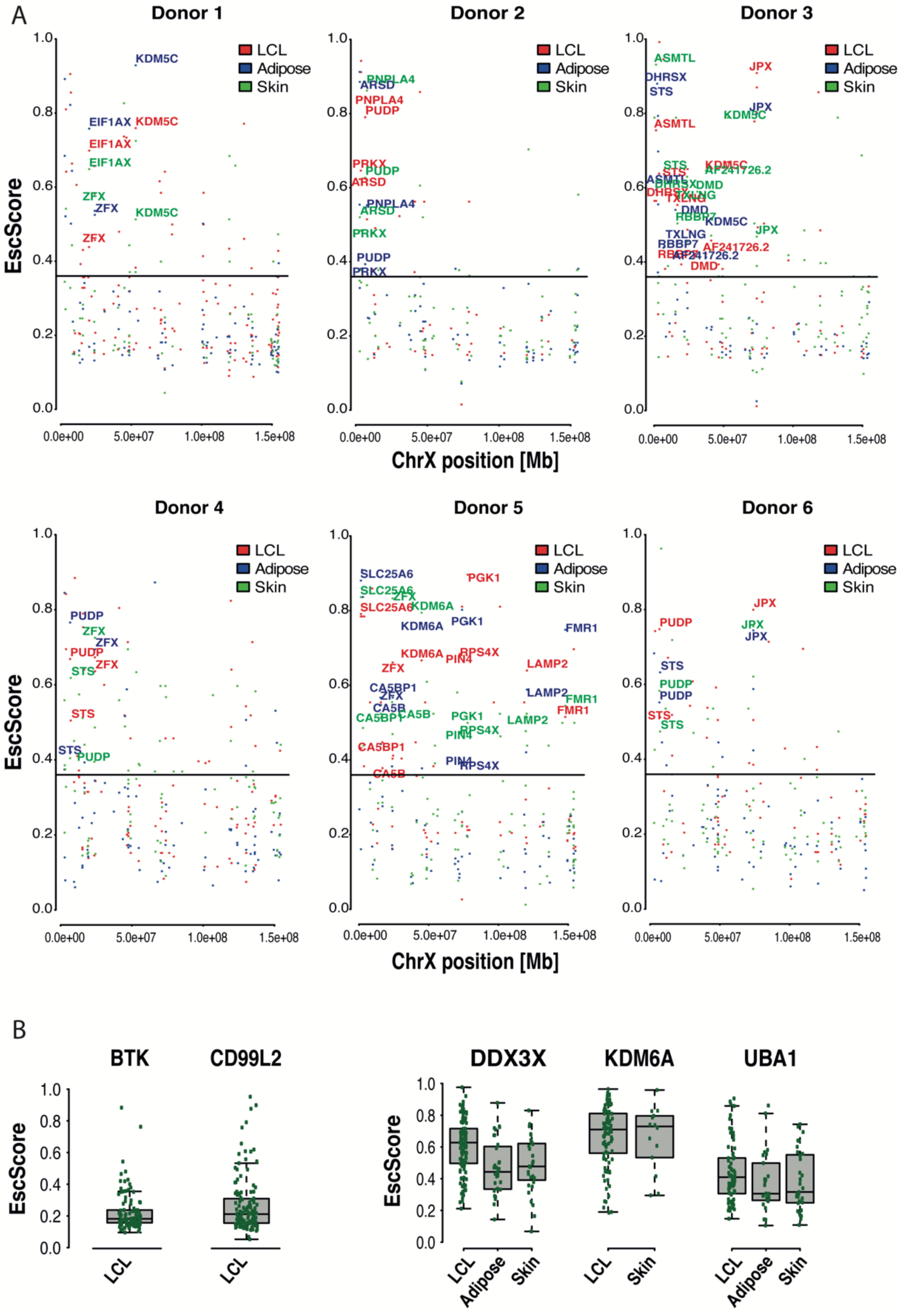
(A) Analysis of intra-individual EscScore variation within each of the 6 female donors exhibiting skewed XCI in all tissues. Each dot shows the gene’s EscScore in each tissue. Genes exhibiting escape (EscScore ≥0.36 in a tissue in a donor) in all 3 tissues within a donor are highlighted in red (LCLs), blue (adipose), green (skin). (B) Representative subset of genes to assess inter-individual variation in EscScore. Each green dot shows the gene’s EscScore in a tissue sample. *BTK* and *CD99L2* show consistent EscScore in LCLs across individuals. *DDX3X, KDM6A* and *UBA1* show inter-individual variability of EscScore in LCLs and a solid tissue. Similar plots for other genes are shown in Fig.S3 and Fig.S4.

To robustly investigate the inter-female diversity in escape, we used a subset of 127 genes escaping XCI in at least 1 tissue, and with EscScore available in at least 10 individuals per tissue. For a given gene, we defined its EscScore to be consistent within a tissue if in at least 80% of individuals the gene’s EscScore lay between ±1 standard deviation from the gene’s average EscScore in the tissue. This strategy revealed both genes with consistent and genes with variable EscScore. We identified 40 genes (∼31% of the interrogated genes) showing consistent EscScore across individuals in at least 1 tissue (Fig.S3). Representative examples are *BTK*, a gene involved in the control of lymphocyte maturation, and *CD99L2*, involved in leucocyte homeostasis. Both genes exhibited consistent EscScore across most donors in LCLs (Fig.2B). A subset of 3 genes (*ARHGAP6, SAT1, RAP2C-AS1*) also exhibited consistency across most donors in both LCLs and a solid tissue (Fig.S3). Genes exhibiting inter-female variability in EscScore in multiple tissues accounted for about 49% (N=62) of the interrogated genes (Fig.S4). Examples are *DDX3X, KDM6A* and *UBA1*, which exhibited variability in LCLs and at least one solid tissue (Fig.2B). Interestingly, inter-female variability occurred more frequently in solid tissues (63% of cases) than LCLs (37% of cases). Altogether, these data are indicative of complex escape patterns. Variable escape across females complements with and may be driven by variable escape across tissues and cells within a female. Inter-female variation has high biomedical relevance as it may underlie predisposition to and manifestation of X-linked traits.

### Escape from X-inactivation exhibits immune cell type-specificity

Females have a higher risk of autoimmune disease than males, and such risk may correlate with increased X-dosage^34,35^. This has raised the hypothesis that XCI escape may contribute to autoimmunity^12,36^. The extent to which escape varies across different immune cells within an individual is not well known. We addressed this question by interrogating 257 X-genes in multiple immune cell types purified from two identical co-twins (Fig.S5). Monocytes, B-cells and T-CD8^+^ cells were available from both co-twins, while T-CD4^+^ and NK-cells from one co-twin. Per each immune cell type, when data were available from both co-twins, we calculated the average gene’s EscScore across the 2 co-twins as a proxy for immune cell type-specific escape. We observed differences between cell types in the average EscScore (Av.EscScore_Monocytes_=0.24; Av.EscScore_B-cells_=0.27; Av.EscScore_T-CD4+_=0.24; Av.EscScore_T-CD8+_=0.28; Av.EscScore_NK-cells_=0.24; KW *P*≤0.01; Fig.3A). These results were consistent when comparison was limited to a subset of 53 genes with EscScore data available in all cell types (Table S8; Av.EscScore_Monocytes_=0.24; Av.EscScore_B-cells_=0.27; Av.EscScore_T-CD4+_=0.26; Av.EscScore_T-CD8+_=0.28; Av.EscScore_NK-cells_=0.24; *P*≤0.01). The incidence of escape varied between cell types, being 15% in monocytes, 20% in B-cells, 22% in T-CD4^+^, 25% in T-CD8^+^, and 28% in NK-cells. Thus, in line with current knowledge, most X-genes are subject to XCI in immune cells. In parallel, our data indicate that escape is heterogeneous across immune cell types, with overall higher incidence in lymphocytes than monocytes. To investigate intra-lineage variation, we compared the EscScore(s) between lymphoid cell types and also found substantial differences (*P*≤0.01), indicating intra-lineage variation. Among the 53 genes with EscScore data available in all cell types (Table S8), we identified 12 genes (*ARSD, PRKX, PUDP, CA5B, AP1S2, ZFX, USP9X, DDX3X, CASK, KDM6A, JPX, DIAPH2*) escaping XCI in at least three immune cell types. *CASK* is a novel candidate escapee. The gene subset *PRKX, ZFX, JPX* and *DIAPH2* escaped XCI in all 5 immune cell types, in line with their behavior as constitutive escapees observed above. For most of these genes, the escape status in immune cells is a novel finding. Interestingly, *KDM6A* exhibited highest EscScore in T-CD8^+^ cells, possibly because of its roles in T-cells control^31^. We identified 9 genes exhibiting escape restricted to one immune cell type, supporting immune cell type-specific factors (Fig.3B). Intriguingly, immune cell type-specific events were restricted to lymphocytes but not monocytes. This might suggest differences between lymphoid and myeloid lineages, and aligns with evidence of increased X-linked biallelic expression in lymphocytes^37^. Altogether, these data indicate that escape varies between immune cell types within an individual. Presumably, this heterogeneity is driven by mechanisms with immune cell type-specific effects.

**FIG.3.**
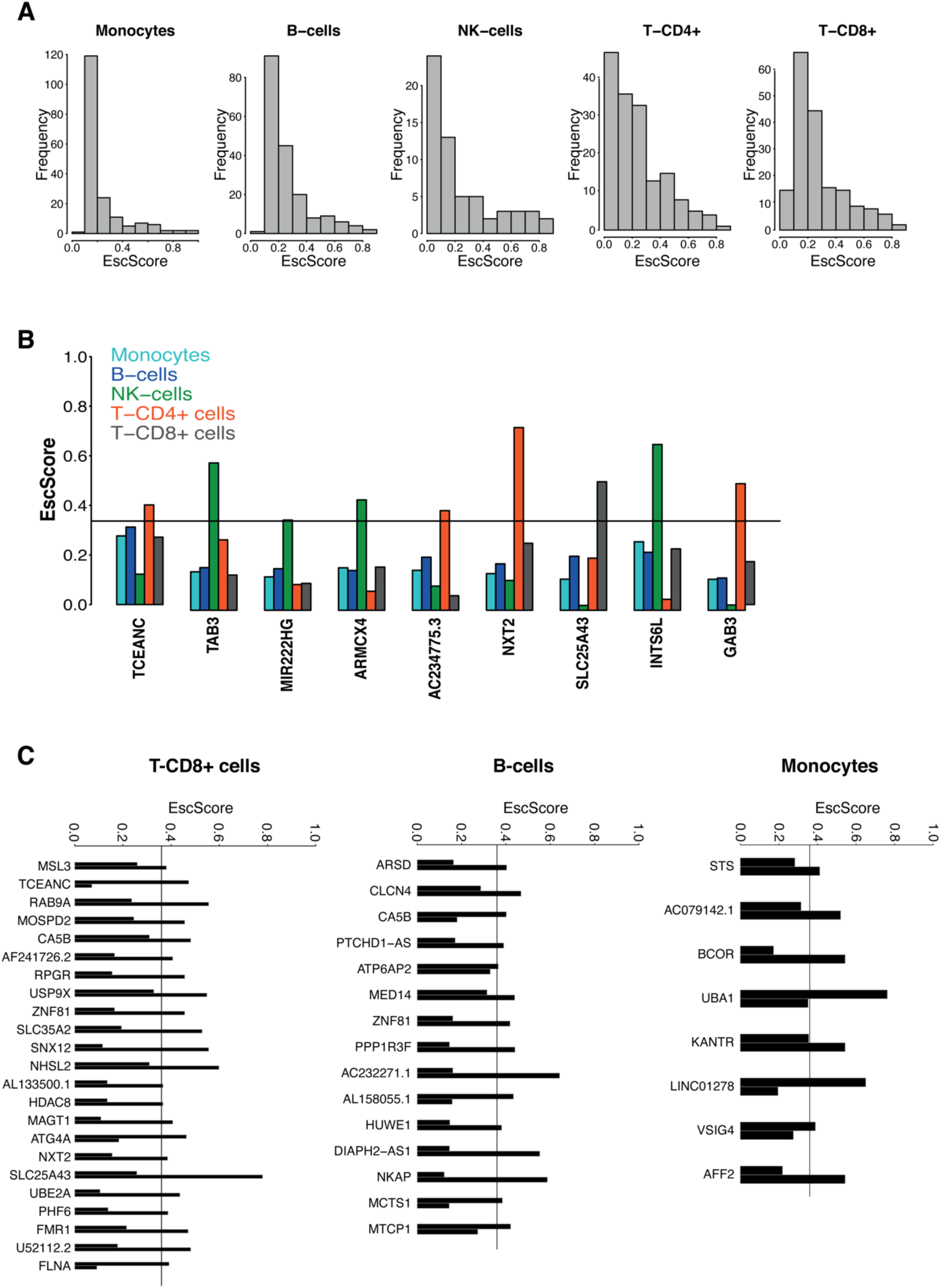
(A) Distribution of EscScore(s) in each immune cell type. (B) Genes exhibiting XCI escape restricted to a single immune cell type. Barplots show the immune cell type-specific gene’s EscScore. (C) Genes exhibiting discordant XCI status (escaping XCI only in one co-twin) between the two MZ co-twins. The two bars shows the gene’s EscScore in an immune cell type (T-CD8^+^ cells, B-cells and monocytes) in the two co-twins.

### Escape from X-inactivation is influenced by heritable and environmental factors

Twin studies are a unique strategy to assess the contribution of genetic factors to complex traits. Using 27 complete twin pairs (17 monozygotic (MZ or identical); 10 dizygotic (DZ or fraternal)), we quantified the concordance in the escape in LCLs between co-twins and compared such concordance between MZ and DZ twins. We correlated the EscScore (using ≥5 genes) between co-twins of each pair (Fig.4A-B), and found that the average correlation across MZ and DZ twins was 0.6 and 0.46, respectively (*ρ*^’^*s* t-test, *P*≤0.05; Fig.4C). These data indicate that MZ share significantly more similar escape than DZ twins. To support this finding, we examined each interrogated gene in a twin pair, and observed higher rates of discordant XCI (gene escaping XCI only in one of the two co-twins) between DZ than MZ twin pairs (Av.Disc.Rate_DZ_=27%; Av.Disc.Rate_MZ_=19%). These data suggest a significant genetic component of escape, in line with previous data on concordance in methylation-based XCI status between MZ twins^16^. In parallel, discordance between MZ twins suggests environmental influences. To gain insights at the cell type level, we next examined the concordance of EscScore in immune cells between two MZ co-twins, and observed significant correlation (*ρ*_monocytes_=0.8; *ρ*_B-cells_=0.68; *ρ*_T-CD8+_=0.6; *P*<1e-10). Genes with discordant XCI status were observed in all three immune cell types, and their prevalence differed between cells ranging from 6.4% in monocytes to 11% in B-cells, and 18% in T-CD8^+^ cells. Interestingly, the genes *CA5B* and *ZNF81* exhibited discordant XCI between the two co-twins in both T-CD8^+^ and B-cells. In all other cases, discordant XCI events concerned distinct gene subsets in distinct immune cell types (Fig 3C). Taken together, our data indicate that genetic and environmental factors may interplay to regulate XCI escape. Variability between immune cell types may also suggest an immune cell type-specific response to environmental influences.

**Fig.4.**
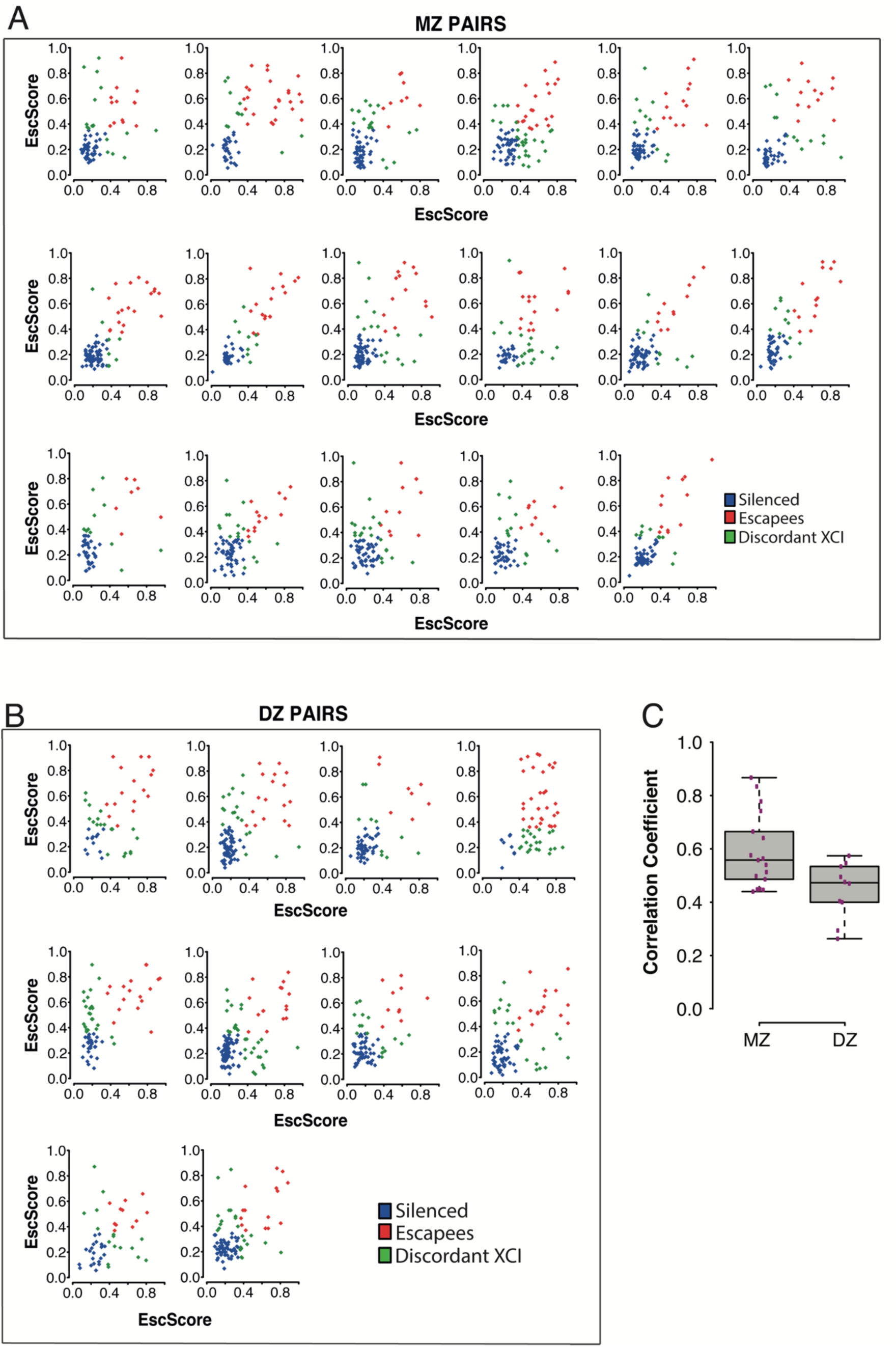
Scatterplots of EscScore(s) of genes (≥5) with data available in both co-twins of a pair. A total of 27 complete twin pairs (both co-twins exhibiting skewed XCI in LCLs) were used. Each dot represents a gene, colored in blue if silenced in both co-twins, red if escaping XCI in both co-twins, and green if exhibiting discordant XCI (escaping only in one co-twin). (A) Monozygotic (MZ) twin pairs (N=17); (B) Dizygotic (DZ) twin pairs (N=10). (C) Boxplots of coefficients of correlation between EscScore(s) in the two co-twins of each pair.

## Discussion

Most knowledge of the extent of escape from X-inactivation in humans is currently based on indirect proxies such as sex comparison, hybrid cell lines, or epigenome analyses^8,11,16,17^. Here, we studied escape in tissues and immune cells using paired transcriptomic and genotype data from nearly 250 female twins from the TwinsUK bioresource^22^. The large sample size and strategy of using tissue samples with skewed XCI^8,9,11,21,38^, enabled us to systematically distinguish silenced from escape genes and identify novel candidate escapees, as predicted^9^. We demonstrate that escape from XCI is a complex phenotype. The incidence of escape varies across tissues, and is higher in solid than blood tissues. Most of our escape calls align with previously annotated XCI statuses, however we identified 54 novel candidate escapees, of which 35 (65%) are protein-coding and 19 (35%) lncRNA genes. We demonstrated via protein network analysis that X-linked protein-coding genes escaping XCI interact with other proteins on a genome-wide scale, and regulate varied biological processes and pathways such as epigenetic changes and hormone signaling. These data indicate that XCI escape has genome-wide effects, in line with recent findings on the effects of XCI changes on global proteome^39^. Thus, de-regulated escape caused by mutations or other disrupting events may have complex phenotypical consequences. Future studies are needed to investigate the functional effects of X-autosomal interactome. Our discovery set includes the PAR1 lncRNA *AL683807*.*1*, which escaped XCI in LCLs, in line with its expression profile in GTEx. The X chromosome is enriched for non-coding RNAs, yet their transcriptional modes and roles are unclear. Due to their unique ability to recruit factors and target a genomic address^1^, lncRNAs play critical roles in genome regulation and health. The study of lncRNAs escaping XCI may reveal novel mechanisms of inter-female phenotypic variation and sexual dimorphism.

We identified genes that constitutively escaped XCI in all tissues, and genes with tissue-specific escape. Co-occurrence of both patterns suggests involvement of tissue-shared and tissue-specific determinants. Presumably, genetic variants (e.g. eQTLs) with constitutive or tissue-specific effects, modulate the escape dosages. We found genes with consistent and with heterogeneous EscScore across individuals. We reasoned that genes with key physiological roles may be subject to shared regulation across females. Examples are *BTK* and *CD99L2*, which, in line with their roles in lymphoid cells, exhibited consistent EscScore in LCLs, but inter-female variability in adipose and skin. Thus, the escape behaviour could depend on the gene’s functional context. The tumorigenesis-related genes *DDX3X* and *KDM6A* both escaped XCI, in line with previous studies^11,25,26^, and showed inter-female variability in all tissues. These genes may underlie sexual dimorphism in cancer^14^, and plausibly their variable escape contributes to inter-female diversity in cancer risk. We found that the histone demethylase *KDM5C* and the transcription factor *ZFX*, escaped in LCLs and solid tissues in the same individual. Other genes manifested more composite behaviour, with escape restricted to either LCLs or solid tissues in an individual, supporting tissue-specificity of escape. Altogether, these patterns highlight the complexity of escape, and anticipate its roles as phenotype modulator.

The X chromosome plays key roles in innate and adaptive immunity^12,27,36,40^. We found that escape differed between immune cell types, with higher incidence in lymphocytes than monocytes. Among 53 genes with data available in all immune cell types, 7% (*PRKX, DIAPH2, JPX, ZFX*) escaped XCI in all cells, in line with their constitutive escape behavior across tissues. For *ZFX*, this aligns with data showing its involvement in networks for X-linked dosage regulation^41^. 17% of genes (*TCEANC, TAB3, MIR222HG, ARMCX4, AC234775*.*3, NTX, SLC25A43, INTS6L, GAB3*) exhibited escape restricted to a single immune cell type which was always of lymphoid lineage. We also found significant variation between lymphoid cells. Altogether, these data indicate an immune cell type-specific propensity to escape. Myeloid and lymphoid lineages are subject to distinct regulation during development. Possibly, cell type-specific factors, such as genetic variants and/or epigenetic marks inherited during differentiation, establish distinct escape dosages across cells within an individual^11^. These data would have multiple biological significance. Firstly, different cell types and possibly single cells, would provide a different contribution to the overall escape dosages in a tissue. This phenomenon could establish the bases for an X-linked transcriptional mosaicism throughout the female’s immune system. Secondly, changes in cell type composition or proportion, which may characterize pathological states^42,43^, may alter the escape which in turn modifies disease risk. The extent to which this phenomenon modulates inter-female variability in risk and expressivity of immunological traits will require future larger studies. Immune cell type-specific escapees could potentially serve as markers of disease-relevant cells, with applications for diagnostic purposes and design of immunotherapy approaches^44-46^.

The extent to which genetics and environment influence escape in humans is unclear. Concordance in methylation-based XCI status between MZ twins supported a dominant model of cis-acting influences^16^. MZ twins share >99% of DNA, age, and multiple environmental traits such as in-utero growth and early life. Variable escape may affect MZ twins differently, leading to different trait expressivity. We found significantly more similar EscScore between MZ than DZ co-twins. Congruently, we found overall higher rates of discordant XCI between DZ than MZ co-twins. These data demonstrate a solid contribution owing to genotype, but also that DNA does not fully explain such concordance patterns. Thus, escape has both heritable and environmental components, in line with current knowledge on complex traits. An interplay between QTLs^47,48^, differential epigenetic control of parental alleles^1^, and gene-environment effects may ultimately modulate the allelic expression of X-genes and propensity to escape. These effects may have cell type- and tissue-specific components, and underlie intra- and inter-female variation across cells and tissues. Our observations on inter-female variability align with population differences in dose compensation^21,49^, supporting the involvement of genetic factors. Cis-acting variation may also model the Xi vs Xa haplotype expression across tissues, explaining intra-individual variation^11^. The identification and functional characterization of such genetic and environmental factors can aid understanding what drives the inter-female variation in trait expressivity and disease risk, and sexual dimorphism. While this is the first multi-tissue transcriptomic study of escape in twins, we acknowledge the need for further strategies to characterize the determinants of XCI escape in humans.

The present study contributes toward a detailed characterization of escape in humans. Given the paradigmatic roles of the X chromosome in epigenetics and clinical genetics, a full understanding of XCI escape has implications on epigenetic research and therapeutics. Therapeutics may include genetic counselling and design of treatments for X-linked conditions. Despite nearly 60 years after Mary Lyon’s landmark intuition on escape, a lot is yet to be learned. Future large-scale studies that combine biomedical records and functional assays will be critical to disentangle the breadth of variability of escape from XCI in humans and characterize its phenotypic impact.

## Materials and Methods

### Sample collection

The study included 856 female twins from the TwinsUK registry^22,50^ who participated in the MuTHER study^47^. Study participants included both monozygotic (MZ) and dizygotic (DZ) twins, aged 38-85 years old (median age = 60). All subjects are of European ancestry. Peripheral blood samples were collected and lymphoblastoid cell lines (LCLs) generated via Epstein-Barr virus (EBV) mediated transformation of B-lymphocytes. Punch biopsies of subcutaneous adipose tissue were taken from a photo-protected area adjacent and inferior to the umbilicus. Skin samples were obtained by dissection from punch biopsies. Adipose and skin samples were weighed and frozen in liquid nitrogen. This project was approved by the research ethics committee at St Thomas’ Hospital (UK). Volunteers received detailed information sheet regarding all aspects of the research, gave informed consent and signed an approved consent form prior to biopsy and to participate in the study.

### DNA sequencing data and variant calling

30X whole genome sequencing (WGS) was carried out at Human Longevity, Inc. (HLI). Details on sample and library preparation, clustering and sequencing have been reported elsewhere^51^. For the purpose of this project, as all individuals were female, the DNA sequencing reads (pre-mapped to the X chromosome by Illumina’s ISIS Analysis Software v.2.5.26.13^52^) were realigned to the GRCh38 X chromosome reference sequence using BWA-MEM in SpeedSeq v0.1.2^53^. Base quality score recalibration (BQSR) was performed in GATK v4.1.6^54^. Following this, DNA variant calling was performed using the gold-standard workflow in GATK v4.1.6^54^. This included implementation of HaplotypeCaller to call germline variants, GenomicsDBImport to create a unified gVCF repository, and GenotypeGVCFs for joint genotyping to produce a multi-sample variant call set. Variants with a VQSLOD (variant quality score odd-ratio) corresponding to a truth sensitivity of <99.9% and with a HWE (Hardy-Weinberg equilibrium) *P-value* <1e-6 were removed. Data quality checks were further performed with VCFtools^55^ to check levels of transition/trasversion ratio. Dataset comprised 621 female samples. For individuals with available RNA-seq but unavailable genotypes, chrX DNAseq data were retrieved from the UK10K project^56^.

### RNA sequencing data

The Illumina TruSeq sample preparation protocol was used to generate cDNA libraries for sequencing. Libraries were sequenced on a Illumina HiSeq2000 machine and 49 bp paired-end reads were generated^50^. Samples that failed library preparation (according to manufacturer’s guidelines) or had less than 10 million reads were discarded. As all individuals were female, for this manuscript RNA-seq reads were aligned to a Y-masked^57^ GRCh38 reference genome using STAR v.2.7.3^58^. Properly paired and uniquely mapped reads with a MAPQ of 255 were retained for further analyses.

### Purified immune cell RNA-sequencing data

Monocytes, B, T-CD4^+^, T-CD8^+^ and NK cells were purified using fluorescence activated cell sorting (FACS) from two monozygotic twins exhibiting skewed XCI patterns in LCLs. Gating strategy for cell sorting is described in Fig.S5. Total RNA was isolated and cDNA libraries for sequencing were generated using the Sureselect sample preparation protocol. Samples were then sequenced with the Illumina HiSeq machine and 126 bp paired-end reads were generated. Adapter and polyA/T nucleotide sequences were trimmed using trim_galore v.0.6.3 and PrinSeq tools v.0.20^59^. RNA-seq reads were aligned to Y-masked^57^ GRCh38 reference genome using STAR v.2.7.3^58^. Properly paired and uniquely mapped reads were retained for further analyses.

### Correction of RNA-sequencing mapping biases

To eliminate mapping biases in RNA-seq, the WASP pipeline for mappability filtering^60^ in STARv2.7.3^58^ was used. In each read overlapping a heterozygous SNP, the allele is flipped to the SNP’s other allele and the read is remapped. Reads that did not remap to the same genomic location are flagged as owing to mapping bias and were discarded.

### Haplotype phasing and gene-level haplotype expression

WGS genomes were read-back phased using recent SHAPEIT2 implementation^61^ that takes advantage of the phase information present in DNA-seq reads. Subsequently, phASER v.0.9.9.4^62,63^ was used for RNAseq-based read-backed phasing and to generate gene-level haplotype expression data. Only reads uniquely mapped and with a base quality ≥10 were used for phasing. Using haplotype expression data, the gene’s ASE in each sample can be calculated as follow:

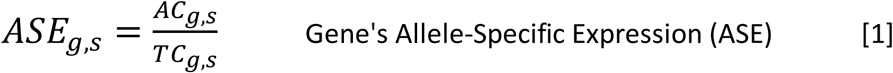

Where, for a biallelic site:

A = haplotype A;

B = haplotype B;

AC_g,s_ = RNAseq allelic count at haplotype A of gene g in sample s;

BC_g,s_ = RNAseq allelic count at haplotype B of gene g in sample s;

TC_g,s_ = Total RNAseq read depth at gene g in sample s;

TC_g,s_ = AC_g,s_ + BC_g,s_.

The gene ASE values range from 0 to 1, with 0 and 1 indicating monoallelic expression and 0.5 indicating completely balanced haplotypic expression. From [1], it follows:

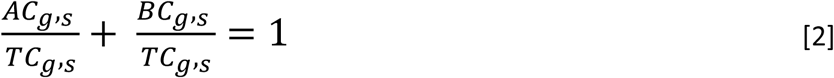

For each gene, the effect size of the allelic imbalance in expression was calculated as allelic fold change (aFC), that is the ratio between the allele with the lower RNAseq count and the allele with the higher RNAseq count, as similarly used in a previous study^64^:

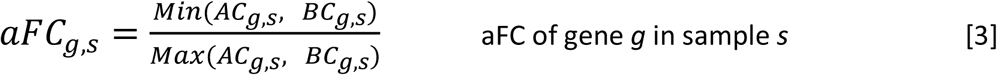

aFC values range from 0 to 1, with 0 indicating monoallelic expression and 1 completely balanced haplotypic expression. For the purpose of our study, a gene’s aFC can be interpreted as Xi/Xa expression ratio. The Xi is assumed to be the allele with the lower RNA-seq count, while the Xa the allele with higher RNA-seq count.

### Quantification of XCI skewing levels

In each sample, the *XIST* allele-specific expression (*XIST*_ASE_) was used as proxy for XCI skewing levels. *XIST* is uniquely expressed from the Xi, and thus the relative expression of parental alleles within *XIST* transcript is representative of XCI skewing in a bulk sample^7,23,24,65,66^. Within each sample, the *XIST*_ASE_ values range from 0 to 1, with 0 or 1 indicating complete inactivation of one parental chromosome (completely skewed XCI, 100:0 XCI ratio), and 0.5 indicating balanced inactivation ratio of the two parental X chromosomes (completely random XCI). To be consistent with previous literature^23,67,68^, we classified samples with *XIST*_ASE_ ≤0.2 or *XIST*_ASE_ ≥0.8 to have skewed XCI, and samples with 0.2< *XIST*_ASE_ < 0.8 to have random XCI. To have an absolute measure of the magnitude of XCI skewing levels in each sample, the degree of XCI skewing (DS) was calculated from the *XIST*_ASE_ calls. DS is defined as the absolute deviation of *XIST*_ASE_ from 0.5, and it has been similarly been used to assess XCI patterns and XCI status of X-genes^21,69,70^. In each sample, DS was calculated as follow:

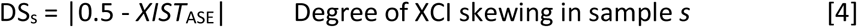

DS is a proxy for the magnitude of the sample’s XCI-skew. DS values range from 0 to 0.5; 0 indicates random XCI and 0.5 completely skewed XCI. Samples with DS ≥0.3 were classified to have skewed XCI; samples with DS<0.3 to have random XCI patterns.

### Quantification of XCI escape

Bulk samples with random XCI patterns confound mono- and bi-allelic X-linked expression as both X-alleles would be, on overall, expressed. Conversely, in skewed samples silenced genes will exhibit monoallelic expression while escape genes biallelic expression^8,11,21^ (Note S1). Only genes with a RNAseq read depth ≥8 reads were used. Furthermore, to increase the confidence that genotypes were truly heterozygous, we considered only genes whose both haplotypes were detected in the RNA-seq data at least once. We reasoned that variations in DS might influence allelic variation in X-linked expression leading to biases in escape measurements across samples. To account for this, we implemented a linear model of the sample’s DS as explanatory variable and the gene’s aFC (as defined above) as response variable using our entire skewed cohort of 166 LCLs, 26 whole-blood, 57 adipose and 64 skin samples. We then derived the aFC’s residuals (referred to as ‘EscScore’), which represent the gene’s aFC adjusted for DS. The EscScore values were subsequently rescaled to the [0,1] range via min-max normalization. Due to low number of skewed whole-blood samples with available data in other tissues, we excluded our whole-blood estimates from further analyses. EscScore 0 and 1 indicate complete monoallelic (silencing) and complete biallelic expression (full escape), respectively. We detected average EscScore of 0.32, 0.36, and 0.37 across LCLs, adipose and skin samples, respectively, whose median is 0.36. These data are consistent with the GTEx XCI survey^11^. Based on these observations, we classified genes exhibiting a median (across ≥3 tissue samples) EscScore≥0.36 as escapees in that tissue, while all others as silenced.

### Protein-protein interaction (PPI) network and gene ontology analyses

Genes with EscScore data available in all three tissues and escaping XCI in at least one tissue were analysed for protein-protein interaction (PPI) with a new genome-wide protein interactome database as reference (www.interactome-atlas.org)^28^. PPI network was imported to STRING v.11^71^, and proteins with at least one direct interaction with our genes and a PPI score (edge confidence) ≥0.4 were selected. PPI network was imported to Cytoscape v.3.8.2^72^ for visualization and gene ontology analyses of Biological Processes and REACTOME Pathways using ClueGO v.2.5.8^73^. A term was considered as significantly enriched when its Bonferroni-corrected *P-value* was ≤0.01.

## Acknowledgements

This study was supported by MRC Project Grant (MR/R023131/1) to K.S.S. The TwinsUK study was funded by the Wellcome Trust and European Community’s Seventh Framework Programme (FP7/2007-2013). The TwinsUK study also receives support from the National Institute for Health Research (NIHR)-funded BioResource, Clinical Research Facility and Biomedical Research Centre based at Guy’s and St Thomas’ NHS Foundation Trust in partnership with King’s College London. This project was enabled through access to the MRC eMedLab Medical Bioinformatics infrastructure, supported by the Medical Research Council [grant number MR/L016311/1]. The authors acknowledge use of the research computing facility at King’s College London, *Rosalind* (https://rosalind.kcl.ac.uk), which is delivered in partnership with the National Institute for Health Research (NIHR) Biomedical Research Centres at South London & Maudsley and Guy’s & St. Thomas’ NHS Foundation Trusts, and part-funded by capital equipment grants from the Maudsley Charity (award 980) and Guy’s & St. Thomas’ Charity (TR130505).

## Authors contributions

A.Z. and K.S.S conceived and designed the project. A.Z. designed and performed bioinformatic analyses. A.L.R. and J.S.E.M contributed data interpretation and discussion. A.V., N.R. and M.F. contributed DNAseq data processing. S.N. performed network and gene ontology analyses and contributed data graphics. R.A.E. performed RNA isolation and FACS experiments. A.Z. and K.S.S. wrote the manuscript. All authors read and approved the manuscript.

## Data Availability

TwinsUK RNAseq data are available from EGA (Accession number: EGAS00001000805). TwinsUK genotypes and phenotypes are available upon application to TwinsUK Data Access Committee (https://twinsuk.ac.uk/resources-for-researchers/access-our-data/). All other data are contained in the manuscript and its supplementary information.

## SUPPLEMENTAL DATA

### Note S1

*XIST* plays essential roles in XCI^1-3^. *XIST* spreads in *cis* from the Xq, triggering an epigenetic silencing of the X designated for inactivation (Xi)^2,4,5^. *XIST* RNA is exclusively expressed from the Xi^3,6,7^, thus the relative expression of parental *XIST* haplotypes can be used to determine the sample’s XCI-skew^8,9^. As XCI occurs at random within each cell, a somatic tissue with random XCI patterns is a mosaic of cells with either parental X silenced. In such a scenario, all X-genes would exhibit biallelic expression, confounding silenced and escape genes^10^. Conversely, in bulk samples with skewed XCI, most expression will be restricted to one haplotype, enabling distinguishment of monoallelic (XCI-silenced genes) from biallelic (escape genes) expression^7,10,11^.

### Note S2

We assessed differences in the incidence of escape between short (Xp) vs long X-arm (Xq). In line with literature^7,12^, we found a higher prevalence of escape on Xp (Fig. S1). This phenomenon has biological explanations as (i) Xp-genes have had Y-paralogs; (ii) the centromere might be a ‘physical barrier’ against the spreading in *cis* of *XIST* from Xq to Xp.

## SUPPLEMENTARY TABLES

**Table S1:**
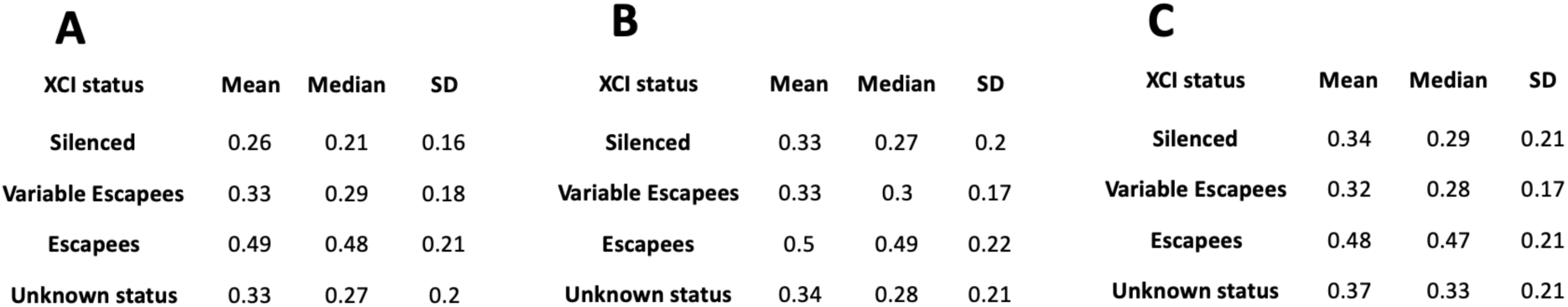
Summary statistics (mean, median, standard deviation) of EscScore values of genes with different annotated XCI status according to the Balaton’s list^13^. A) LCLs; B) adipose; C) skin. Each statistics is computed across ≥3 tissue samples.

**Table S2:** Statistical comparison between the EscScore(s) of different gene categories previously annotated^13^. Escapees refer to genes annotated as fully or mostly escaping XCI. For each interrogated gene, the median EscScore value across ≥ 3 tissue samples was used for comparison. Tables report the p-value of the Wilcoxon test between two gene categories A) LCLs; B) adipose; C) Skin.

| **A** |  |  |  | **B** |  |  |  | **C** |  |  |  |
| --- | --- | --- | --- | --- | --- | --- | --- | --- | --- | --- | --- |
| XCI status | Silenced | Variably Escapees | Escapees | XCI status | Silenced | Variably Escapees | Escapees | XCI status | Silenced | Variably Escapees | Escapees |
| Silenced | * | 0.1 | 9e-24 | Silenced | * | 0.3 | 4e-14 | Silenced | * | 0.7 | 1e-14 |
| Variably Escapees | 0.1 | * | 9e-8 | Variably Escapees | 0.3 | * | 1e-4 | Variably Escapees | 0.7 | * | 2e-4 |
| Escapees | 9e-24 | 9e-8 | * | Escapees | 4e-14 | 1e-4 | * | Escapees | 1e-14 | 2e-4 | * |

**Tables S3, S4, S5, S6, S8 are described below and provided as separate files:**

**Table_S3**: X-linked genes (N=159) exhibiting escape in at least one of the three studied tissues (LCLs, adipose, skin) in our dataset. The table lists the gene’s EscScore in each tissue, computed as the median EscScore across ≥3 samples.

**Table_S4:** X-linked genes (N=215) with EscScore available in all three studied tissues (LCLs, adipose, skin). The table lists the gene’s EscScore in each tissue, computed as the median EscScore across ≥3 samples.

**Table_S5:** X-linked genes (N=51) escaping XCI in only one of the three studied tissues (LCLs, adipose, skin). This is a subset of Table_S4. Column 2 indicates whether the escape status of the gene was either known (previously reported) or is a novel call.

**Table_S6:** Results from ClueGO^14^ enrichment analysis of genes in the PPI network.

**Table_S7.** Fraction of X-linked genes exhibiting escape (EscScore ≥0.36 in a tissue in a donor) in 1, 2 or all 3 studied tissues in each of the 6 female donors exhibiting skewed XCI in all three studied tissues (LCLs, adipose, skin).

| **DONOR** | % escape in 1 tissue | % escape in 2 tissues | % escape in 3 tissues |
| --- | --- | --- | --- |
| **Donor 1** | 17.8% | 5.6% | 1.6% |
| **Donor 2** | 17.4% | 3.5% | 3.5% |
| **Donor 3** | 26.6% | 7.8% | 7% |
| **Donor 4** | 35% | 6% | 2% |
| **Donor 5** | 27.5% | 3.8% | 7.6% |
| **Donor 6** | 30% | 2.6% | 2.6% |

**Table_S8:** X-linked genes (N=53) with EscScore available in all five studied immune cell types (Monocytes, B-cells, T-CD4^+^ cells, T-CD8^+^ cells, NK-cells). The table lists the genes’ EscScore in each immune cell type.

## SUPPLEMENTARY FIGURES

**Fig.S1:**
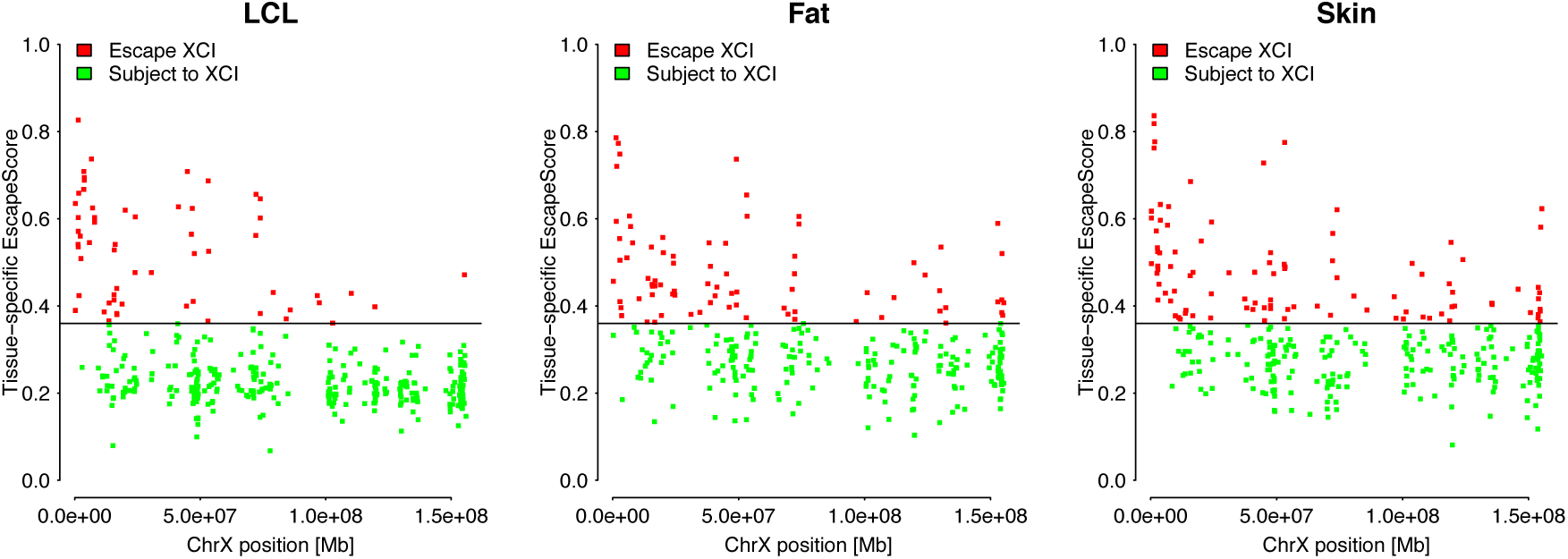
Relationship between the gene’s tissue-specific EscScore and gene position on chrX (GRCh38). Each dot represents a gene. Red and green dots are escapee and silenced genes, respectively.

**Fig.S2:**
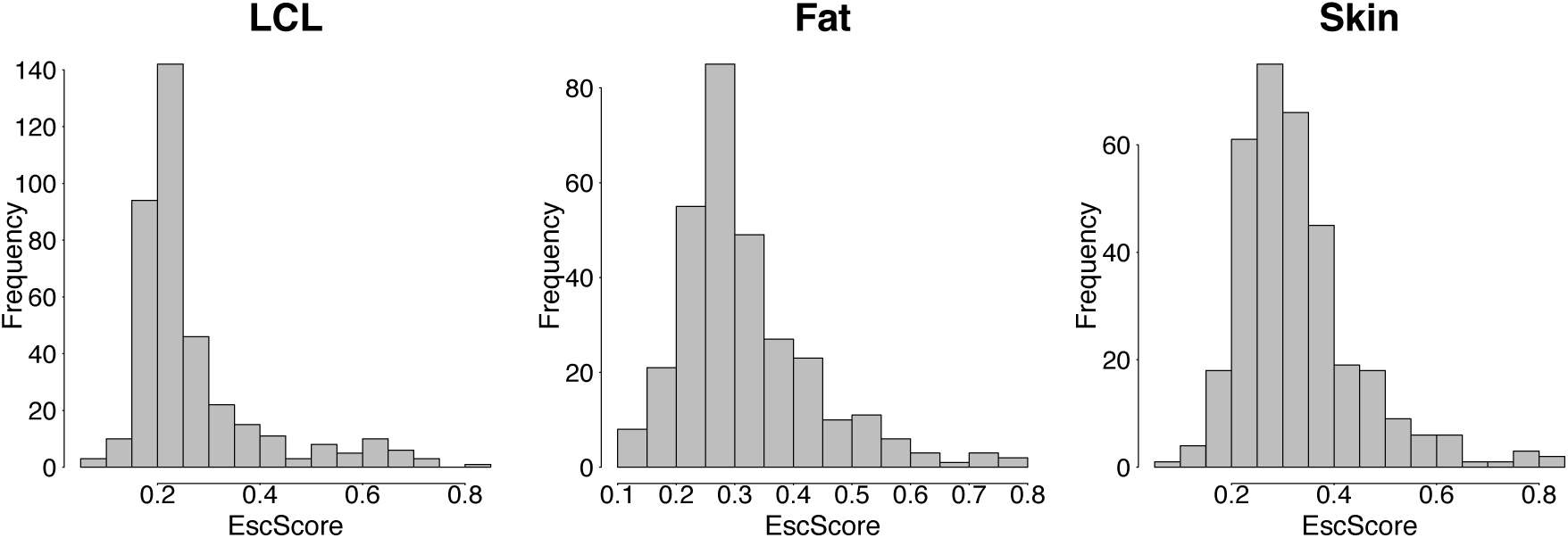
Distribution of median EscScore values in each of the three studied tissues (LCLs, adipose, skin). Median values were calculated per gene across ≥3 skewed tissue samples.

**Fig.S5:**
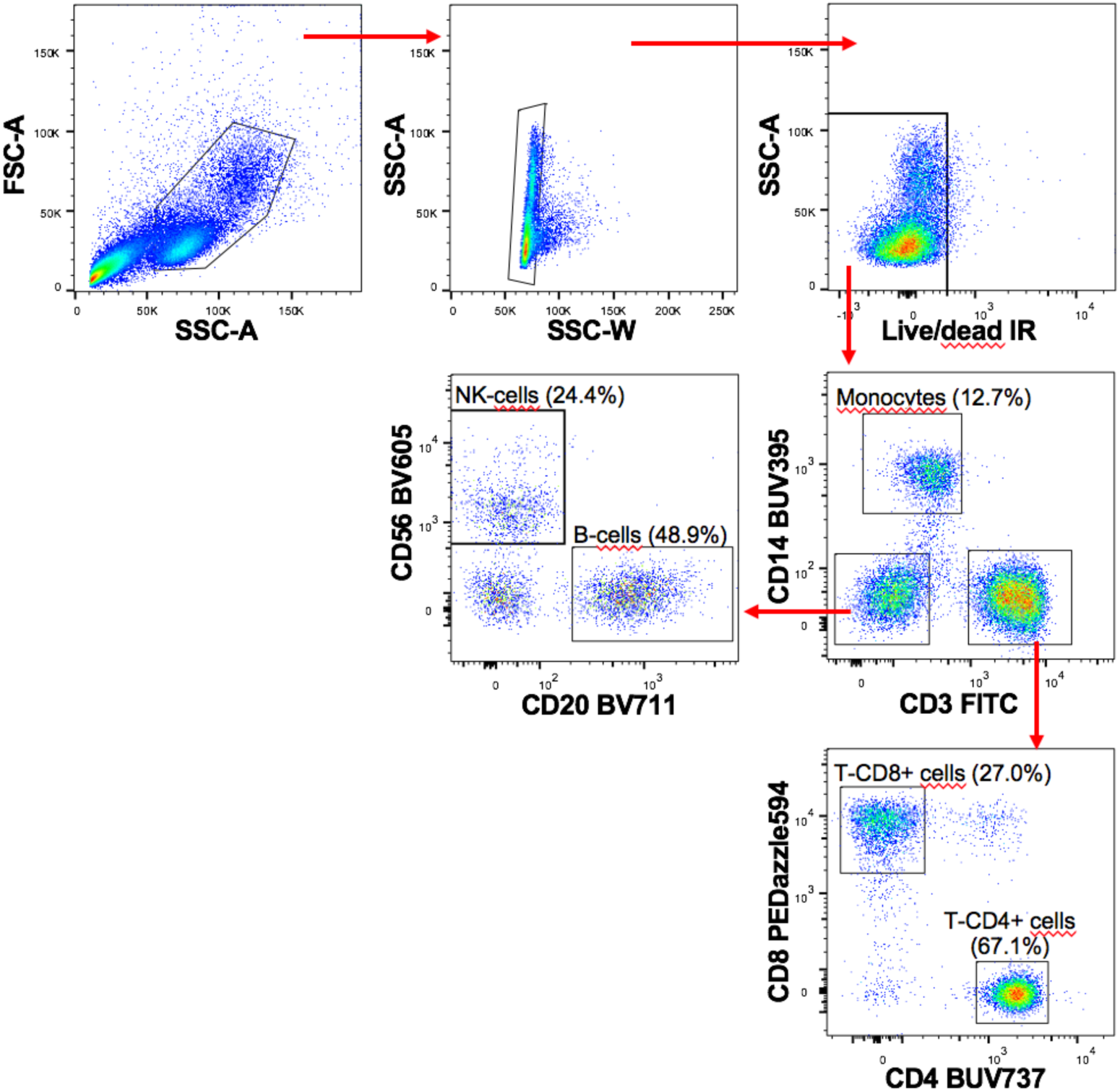
Gating strategy for immune cell sorting. Gating strategy used to sort monocytes (CD14+), B (CD14-, CD3-, CD56-, CD20+), NK (CD14-, CD3-, CD20-, CD56+), T-CD4+ (CD14, CD3+, CD8-, CD4+) and T-CD8+ cells (CD14-, CD3+, CD4-, CD8+) from freshly isolated PBMCs from 2 monozygotic twins exhibiting skewed XCI in LCLs.

**Fig_S3.pdf [Pages 26-27]:**
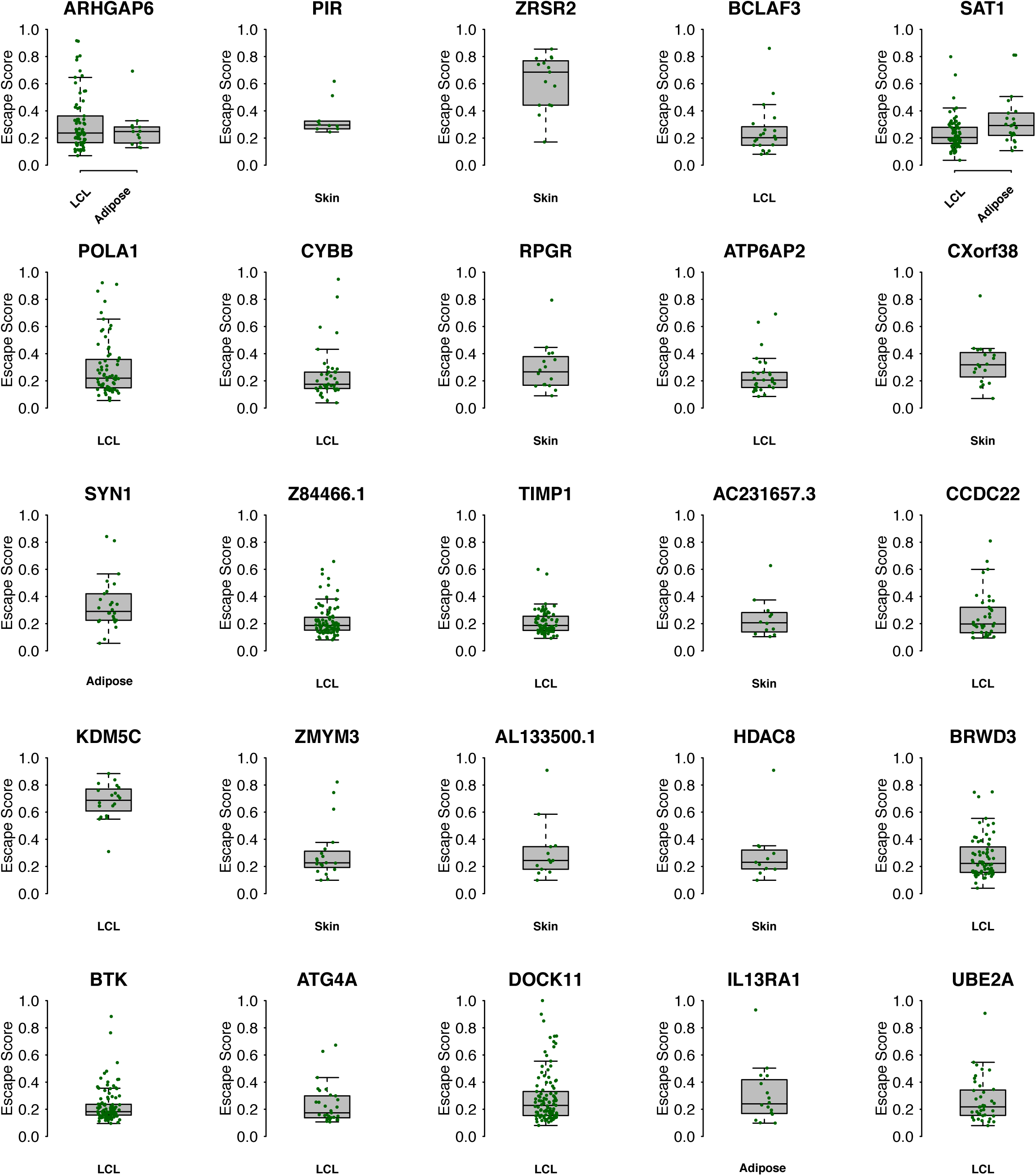
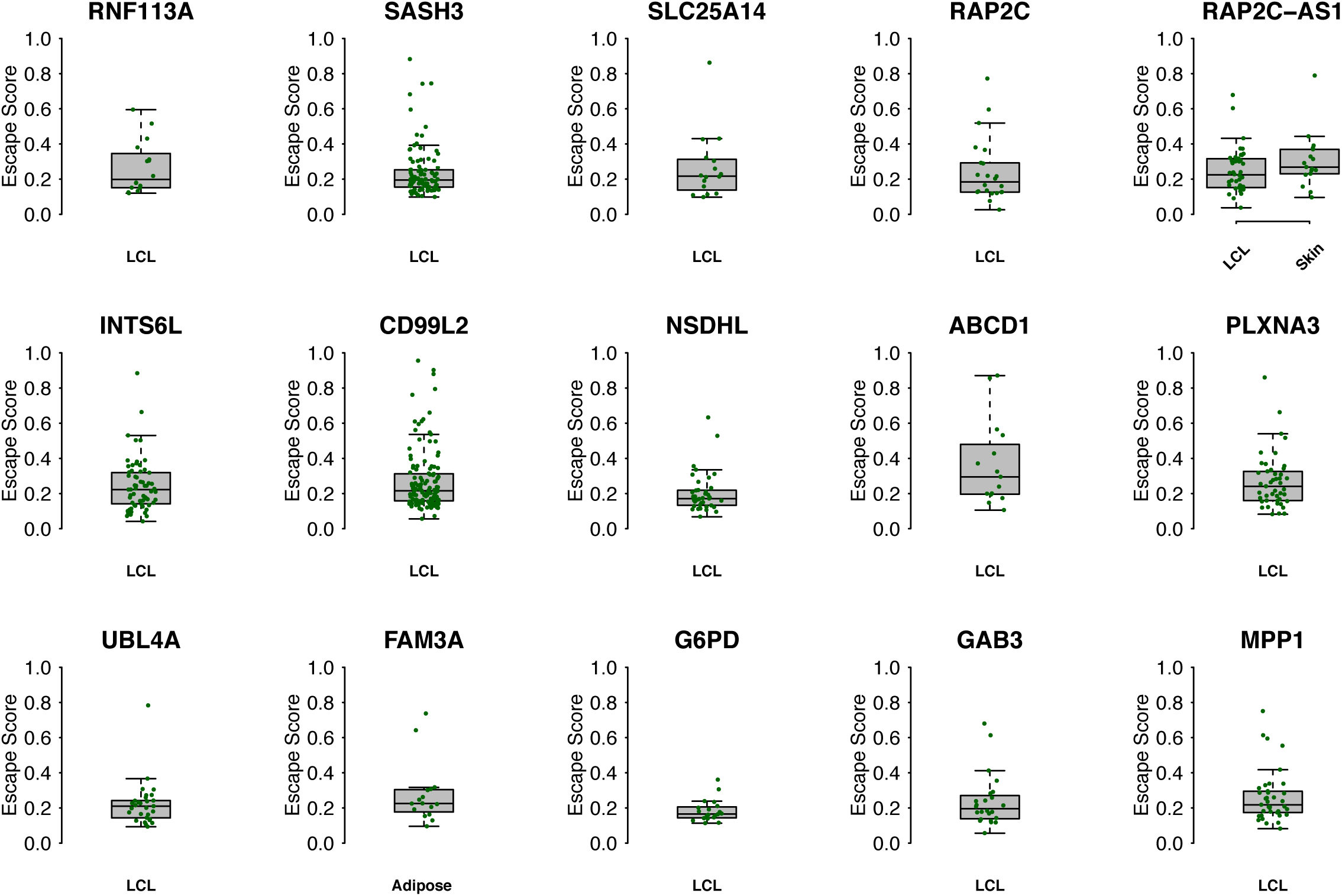
Boxplot of gene’s EscScore in each skewed sample (≥10 tissue samples used for this analysis). Plotted are genes classified to have consistent EscScore across individuals. Each green dot is an individual.

**Fig_S4.pdf [Pages 28-32]:**
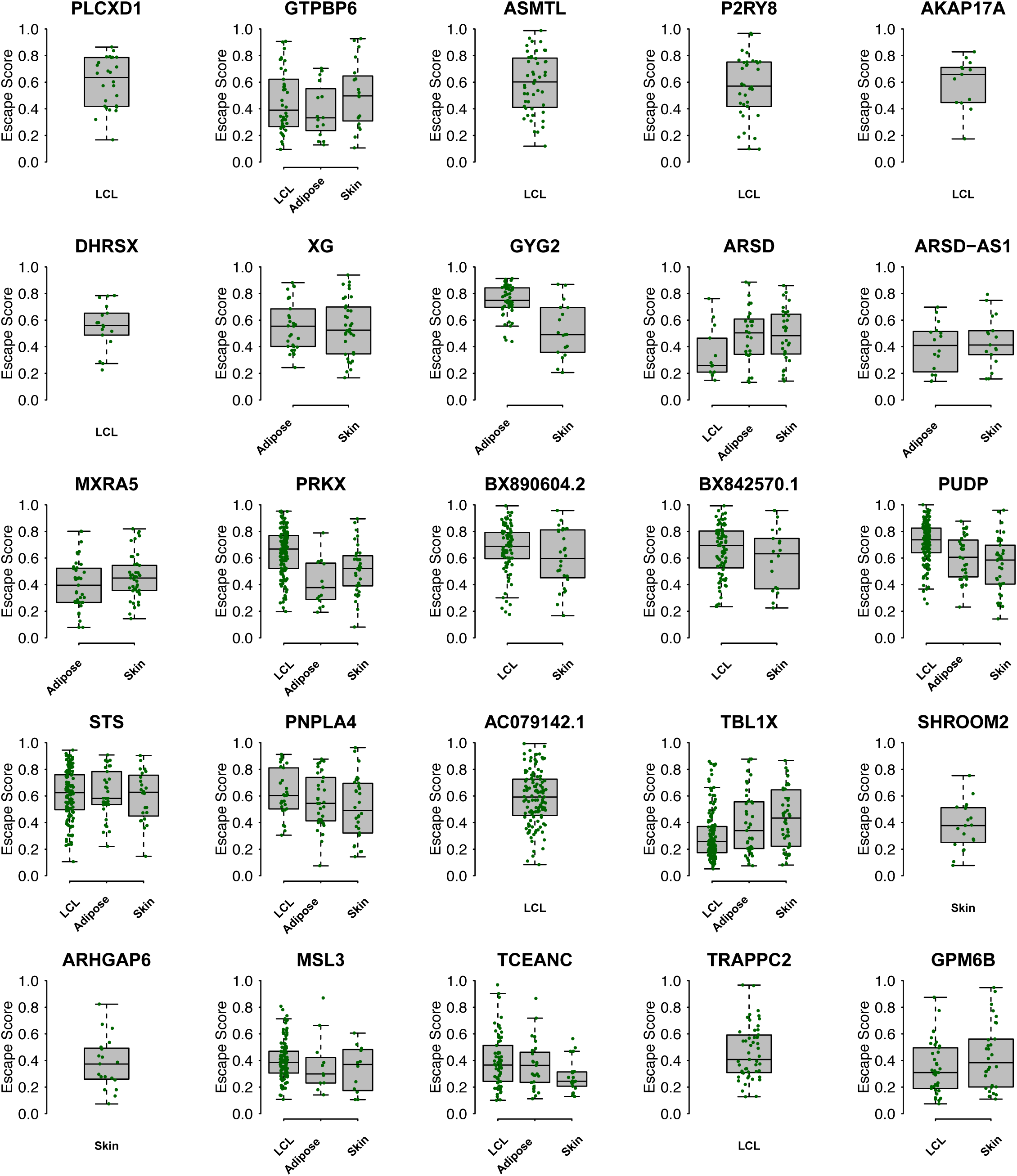
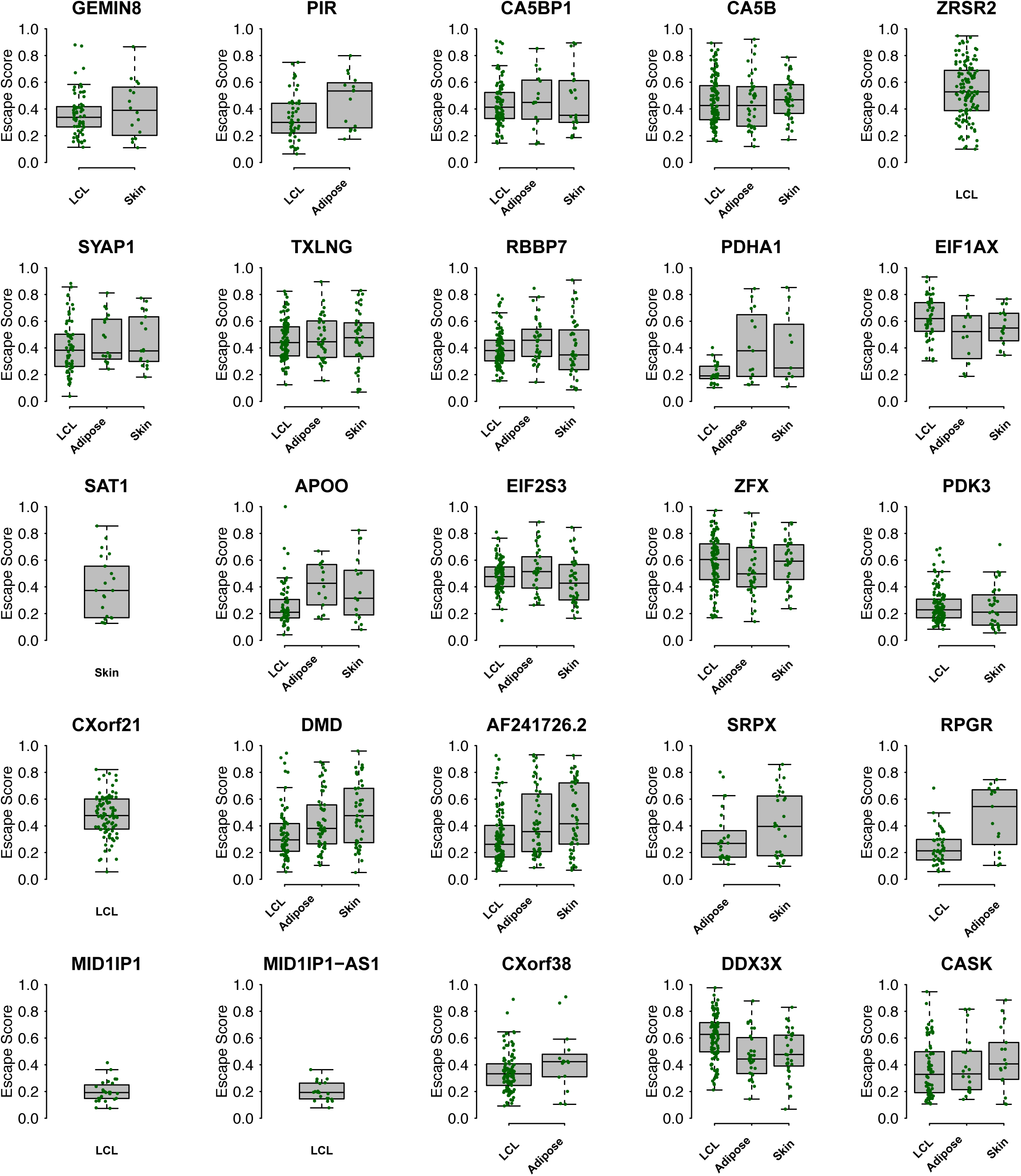
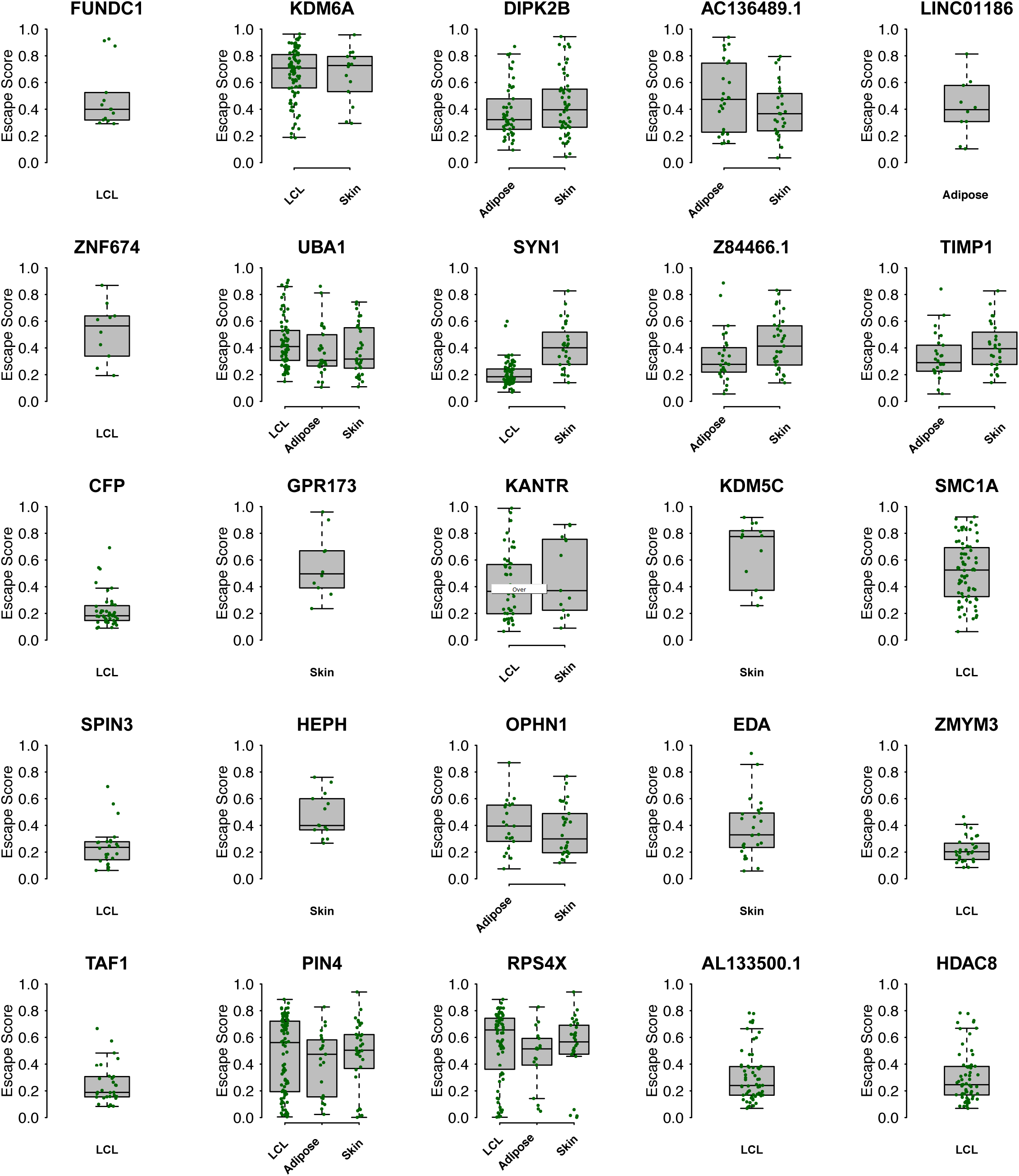
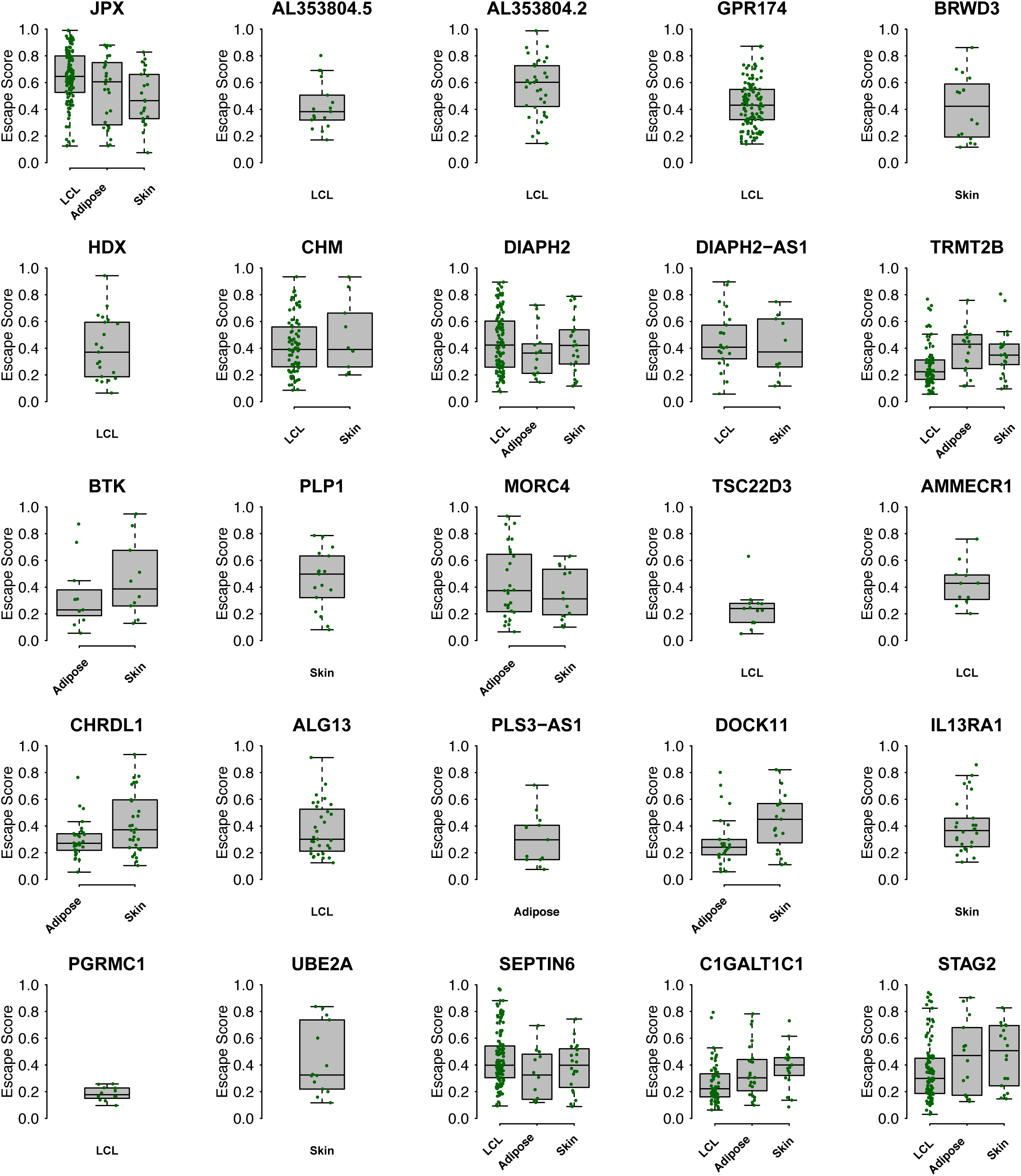
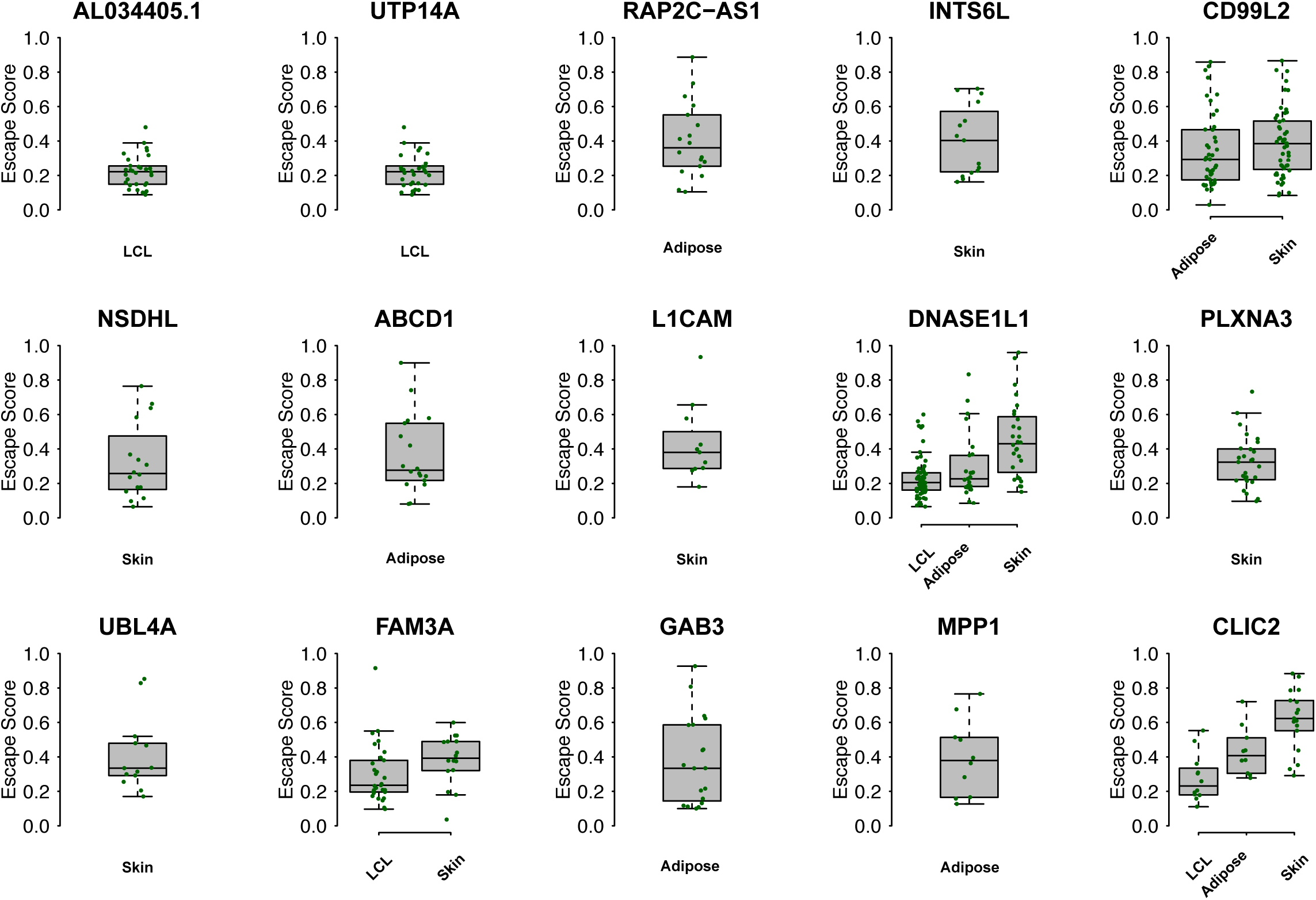
Boxplot of gene’s EscScore in each skewed sample (≥10 tissue samples used for this analysis). Plotted are genes classified to have variable EscScore across individuals. Each green dot is an individual.

